# Trace Amines are Essential Metabolites for the Autocrine Regulation of *β*-Cell Signaling and Insulin Secretion

**DOI:** 10.1101/2024.03.21.585773

**Authors:** Sebastian Hauke, Kaya Keutler, Aurelien Laguerre, Mireia A. Carbo, Jona Rada, David Grandy, Dmytro A. Yushchenko, Carsten Schultz

**Author notes:** authors contributed equally to this work (Sebastian Hauke, Kaya Keutler). To whom correspondence should be addressed: C. Schultz. S.H. present address: AbbVie Deutschland GmbH & Co. KG. NBE analytical R&B, Knollstraße, 67061 Ludwigshafen. D.A.Y. Present address: Miltenyi Biotec B.V. & Co. KG, Friedrich-Ebert-Straße 68, 51429 Bergisch Gladbach, Germany.

## Abstract

Secretion of insulin in response to extracellular stimuli, such as elevated glucose levels and small molecules that act on G-protein coupled receptors (GPCRs), is the hallmark of *β*-cell physiology. Trace amines (TAs) are small aromatic metabolites that were identified as low-abundant ligands of the trace amine-associated receptor 1 (TAAR1) in the central nervous system (CNS), a GPCR that is also expressed by pancreatic *β*-cells. In the present work, we identify TAs as essential autocrine signaling factors for *β*-cell activity and insulin secretion. We find that *β*-cells are producing TAs in significant amounts and that the modulation of endogenous TA levels by the selective inhibition of TA biosynthetic pathways directly translated into changes of oscillations of the intracellular Ca^2+^ concentration ([Ca^2+^]_i_ oscillations) and insulin secretion. Selective TAAR1 agonists or inhibitors of monoamine oxidases increased [Ca^2+^]_i_ oscillations and insulin secretion. Opposite effects were mediated by selective TAAR1 antagonists, by recombinant monoamine oxidase action and by the inhibition of amino acid decarboxylase. As the modulation of TA biochemical pathways immediately translated into changes of [Ca^2+^]_i_ oscillations, we inferred high metabolic turnover rates of TAs and autocrine feedback. We found that psychotropic drugs modulate [Ca^2+^]_i_ oscillations and insulin secretion, either directly acting on TAAR1 or by altering endogenous TA levels. Our combined data support the hypothesis of TAs as essential autocrine signaling factors for *β*-cell activity and insulin secretion as well as TAAR1 as an important mediator of amine-modulated insulin secretion.

## INTRODUCTION

Insulin secretion is a synchronized multi-cellular process. Coordinated oscillations of the intracellular free Ca^2+^ concentration ([Ca^2+^]_i_ oscillations) along with periodic secretory bursts of insulin are ensured by the integration of the pancreatic *β-*cell population into functional syncytia within islets of Langerhans [1–5]. Direct gap-junctional coupling of *β-*cells by connexins has been described as the basic mechanism for *β-*cell synchronization [6–9]. However, loss of functional gap junctions or dissociation of pancreatic islets into single *β-*cells was shown to compromise, but not to abolish synchronized [Ca^2+^]_i_ oscillations and pulsatile insulin release [10,11]. In addition to gap junctions [9], the role of soluble para- and autocrine signaling factors in the extracellular space of pancreatic islets has come into focus for *β*-cell synchronization and insulin secretion [12–14]. Pancreatic *β*-cells express a multitude of GPCRs [15], such as the fatty acid (FA) receptor GPR40 that was found to mediate the majority of the effects that long-chain FAs have on *β*-cell activity [16]. In our recent work, we demonstrated the essential role of cellular FAs and ATP for [Ca^2+^]_i_ oscillations and insulin secretion [12,17]. We found that ATP and endogenous FAs do not only stimulate *β*-cells, but are secreted by these cells and are required for basal *β*-cellular activity and insulin secretion [12,17]. Other diffusible autocrine ligands of GPCRs include NO [18,19], CO [20], Zn^2+^ [21], neuropeptide Y [22], glutamate [23] and γ-aminobutyric acid [24].

In 2001, the ‘trace-amine associated receptor 1’ (TAAR1) was de-orphanized as a receptor for trace amines (TAs) that are low abundant aromatic metabolites [25,26]. TAs are generated from amino acids by aromatic L-amino acid decarboxylase (AADC) and are inactivated by monoamine oxidase (MAO) action [27,28]. Since its discovery, research has mainly focused on the neuromodulatory functions of TAAR1, making it a novel target for the treatment of psychiatric disorders [29–32]. TAs have been implicated in the modulation of aminergic neurotransmission in the CNS [33]. Deviations of TA levels have been linked to common psychiatric disorders, such as major depression [34,35], schizophrenia [36], Parkinson’s disease and anxiety states [30,37]. Therefore, a series of potent synthetic full (*e.g.* RO5166017, RO5256390) and partial TAAR1 agonists (*e.g.* RO5073012, RO5203648 and RO5263397) were developed for the modulation of TAAR1 activity with high selectivity over the adrenergic α2 receptor [29–31,38]. RO5212773 (EPPTB) is currently the only commercially available TAAR1 antagonist [33]. Based on homology modeling, Tan *et al.* predicted lead antagonist properties for rat TAAR1 to develop the potent TAAR1 antagonist ET-92 [39].

Much less is known about the effects of TAAR1 ligands in the periphery although TAAR1 expression has recently been reported in organs involved in gastric function and nutrient absorption, such as the duodenum, stomach and the pancreas [29,40–42]. TAAR1 has been found amongst the most highly expressed GPCRs in *β-*cells of human and mouse pancreatic islets, as well as in some pancreatic cell lines [40,42]. Recently, TAAR1-mediated downstream signaling and its effects on glucose-stimulated insulin secretion (GSIS) as well as *β-*cell proliferation were investigated in the rodent *β-*cell lines INS-1 and MIN6 [43]. Through coupling to G_αs_, stimulation of TAAR1 has been reported to trigger cAMP-mediated Ca^2+^ influx, PKA-mediated CREB phosphorylation along with upregulation of insulin receptor substrate-2 (Irs-2) expression [43]. In line with this, selective synthetic and endogenous TAAR1 agonists were shown to potentiate GSIS from rat INS-1E cells, from isolated human islets and in C57BL/6 mice, along with significantly lowered blood glucose levels [40,42]. These studies were the first to describe TAAR1 as an integrator of metabolic control over insulin secretion. Despite data demonstrating TAAR1 expression in the pancreas, the identity of endogenous ligands of pancreatic TAAR1 and the role of receptor stimulation by endogenous ligands for individual *β*-cell activity and insulin secretion is yet to be determined.

In the present study, we demonstrate that TAs are produced and secreted by pancreatic *β-*cells. We identify these endogenous TAs as essential autocrine signaling factors for *β-*cell activity and insulin secretion. We show that *β-*cell activity can therefore be modulated by drug-mediated reduction or increase of endogenous TA levels. Our approach to subordinate *β-*cell activity and insulin secretion to the activation of the yet underestimated TAAR1 in the pancreas by small aromatic metabolites contributes to a more detailed understanding of the fundamental regulation of *β-*cell activity and insulin secretion.

## RESEARCH DESIGN AND METHODS

### Reagents

Dulbeccós Modified Eagle Medium (DMEM) was obtained from Gibco (Carlsbad, Ca). *β*-Mercaptoethanol (50 mM in PBS) was ordered from PAN Biotech (Aidenbach, Germany). Minimal medium for culturing cells in the presence of isotope-labeled amino acids was delivered by Cell Culture Technologies (Gravesano Ticino, Switzerland). Collagenase (from *Clostridium histolyticum*) was obtained from Nordmark Biochemicals (Uetersen, Germany). Histopague 1083 (density: 1.083 g/mL) and Histopague 1119 (density: 1.119 g/mL) were purchased from Sigma-Aldrich. Stable isotope-labelled amino acids and trace amines were purchased from Sigma Aldrich, CDN Isotopes (Quebec, Canada) and Santa Cruz Biotechnology (Huissen, The Netherlands). Caged phenylethylamine was prepared as described in the Supplemental Data. Solvents of the highest analytical grade were provided by Biosolve (Valkenswaard, The Netherlands), Honeywell (Morristown, US) and Sigma-Aldrich. All other commercially available chemicals and enzymes were ordered from Sigma-Aldrich. All anesthetics used were purchased and stored in compliance with the German Narcotics Act (BtMG).

### Unit definitions of applied recombinant enzymes

According to the supplier, Sigma Aldrich: *Monoamine oxidase B* (from baculovirus infected insect cells, EC number: 1.4.3.4): One unit deaminates 1.0 nmole of Kynuramine per minute at pH 7.4 and 37 °C. Before application on cells, monoamine oxidase was re-buffered by HEPES-based buffer as applied in imaging experiments (see below).

### Culturing MIN6 cells

MIN6 cells [44] were cultured in a humidified atmosphere at 37 °C and 8% CO_2_. The culture medium DMEM contained 4.5 g/L glucose and was supplemented with FBS (15%) and *β*-mercaptoethanol (70 µM). The medium was sterile-filtered (Millex GV, 0.22 µm) and used within one week after preparation. For imaging, MIN6 cells were seeded into 8-well LabTek microscope dishes (155411 Thermo Scientific) or 40 mm coverslips (Menzel Gläser, Braunschweig, Germany). For *in vitro* assays, MIN6 cells were seeded into 0 60 mm or 0 35 mm dishes (Nunc delta surface, cat# 150288 / cat# 153066; Roskilde, Denmark) to form pseudoislets within 5 days after seeding. MIN6 cells were a kind gift from Prof. T. Miyazaki and Prof. K. Yamamura, Kumamoto University Medical School, Japan, and were used from passages 27 - 36. MIN6 cells were cultured in the presence of stable isotope-labeled amino acids. Minimal medium, depleted of L-tyrosine, L-tryptophan and L-phenylalanine (Cell Culture Technologies (Switzerland)) was supplemented with stable isotope-labeled (^15^N) amino acids L-tyrosine, L-tryptophan and L-phenylalanine (0.4 mM, 0.08 mM, and 0.4 mM). The pH was adjusted to 7.0. Before use, the medium was supplemented with FBS (15%), *β*-mercaptoethanol (70 μM), and was sterile-filtered.

### Isolation of mouse primary *β*-cells

Female Ctrl: CD1 (ICR, outbred) mice (supplied by Charles River Laboratories, cat# CD1S1FE07W) served as donors for primary *β*-cells that were isolated as previously described [45]. In short, mice were anesthetized in a CO_2_ atmosphere and sacrificed by cervical dislocation. A collagenase solution (1 mg/mL) was injected into the pancreatic duct, followed by extraction of the pancreas from the animal and digestion at 37 °C for 10 min. A Histopague gradient (1.083 and 1.119 g/mL) allowed for the isolation of pancreatic islets via density gradient centrifugation. Islets were incubated in RPMI medium (supplemented with 10% FCS, 100 U/mL penicillin and 100 mg/mL streptomycin) for 24 h. Trypsin digestion (5 min, 37°C) dissociated islets into single *β*-cells that were seeded into LabTeks, pre-coated with poly-L-lysine. Animals were housed in the EMBL animal facilities under veterinarian supervision and the guidelines of the European Commission, revised directive 2010/63/EU and AVMA guidelines 2007.

### Mouse insulin ELISA

The quantification of insulin secretion was based on an enzyme-linked immunosorbent assay (ELISA) in 96-well format. For the quantification of insulin release from MIN6 or mouse primary *β*-cells, a mouse insulin ELISA kit (serial number: 10-1247, Mercodia AB, Uppsala, Sweden) was applied. Insulin release from INS-1 cells was measured using a rat insulin ELISA kit (serial number: 10-1250, Mercodia AB, Uppsala, Sweden). Experiments for the determination of insulin secretion were performed in quadruplicate per condition, following the instructions of the supplier. Insulin levels were normalized to the protein content of MIN6 or mouse primary *β*-cells, as determined by a BCA assay (Pierce^TM^ BCA Protein assay kit, Thermo Scientific, Rockford, IL, USA). Artifacts of BSA on insulin measurements were corrected according to Fig. A23 (appendix).

### Confocal laser scanning microscopy

Live cell imaging was performed on a FluoView1200 (Olympus IX83) confocal laser scanning microscope, equipped up with an environment box (custom-made at EMBL) to allow imaging at 37°C and 5% CO_2_. Olympus 60x Plan-APON (NA 1.4, oil) or 20x UPLS APO (NA 0.75, air) objectives and FluoView software (version 4.2) were applied. Images were acquired using a Hamamatsu C9100-50 EM CCD camera. A 488 nm laser line (120 mW/cm^2^, 2.5 %) in combination with a 525/50 emission mirror was used to image the green channel. A 559 nm laser line (120 mW/cm^2^, 2.0%) and a 643/50 emission filter were used for red channel recordings. A pulsed UV laser line (375 nm, 10 MHz) was applied for uncaging experiments on live cells. Uncaging was performed in the entire field of view for eight frames (frame time of uncaging: 3.2 s/fame). A dual-scanner set-up allowed for image acquisition of cellular responses during UV stimulation. For monitoring changes of [Ca^2+^]_i_ in response to external stimuli, cells were incubated with the acetoxymethyl ester of the Ca^2+^ indicator Fluo-4 (Life Technologies, Eugene, OR), 5 µM in DMEM (1 g/L glucose) for 25 min at 37 °C and 5 % CO_2_. The frame time was set to 3.9 s, with images acquired in 4 s intervals. For imaging, MIN6 cells were grown as pseudo islets of ∼70 % confluence. Imaging experiments were performed in HEPES buffer (in mM: 115 NaCl, 1.2 CaCl_2_, 1.2 MgCl_2_, 1.2 K_2_HPO_4_ and 20 HEPES, pH 7.4).

### Imaging of intracellular cAMP levels using an EPAC-based sensor

For monitoring changes of intracellular cAMP levels upon TAAR1 stimulation, MIN6 cells were transiently transfected with Epac^3rd^ [mTurquoise-Epac(CD,delDEP)-^cp173^Venus-Venus] [46]. For this, we used Lipofectamine 3000, following the guidelines of the supplier. Stimuli were applied to cells in 25 µM final concentrations in the presence of 11 mM glucose. Imaging was performed on an Olympus FV1200 microscope, using an Olympus 60x Plan-APON (NA 1.4, oil) objective in a sealed chamber (EMBL incubation box) at 37°C and 5% CO_2_. Settings: Frame time: 5 s with 10.0 µs/pixel (640x640 pixels), laser power 1.5 %, pinhole: 600 µm. Epac^3rd^ was excited at λ = 405 nm. Optical filters were applied for mTurquoise (λ = 450 - 490 nm) and for Venus (λ = 510 - 560 nm) emission. FRET values are presented as the ratio between donor and acceptor signals. The FRET ratio was normalized to a value of one at the time point immediately before adding the stimulus. At the end of each experiment, a mix of the adenylate cyclase activator forskolin and the phosphodiesterase inhibitor 3-isobutyl-1-methylxanthine (IBMX) (50 µM, each) was added to MIN6 cells to stimulate maximum cAMP responses.

### Analysis of imaging data

The open-access software tool Fiji [47] was applied for the extraction of fluorescence intensities from individual cells. Following background subtraction, intensities were calculated relative to the maximum detected fluorescence intensity (F/F_0_). Representative traces of cells within a field of view were averaged or numbers of detected high-intensity [Ca^2+^]_i_ events per 60 s interval were determined. The height of each [Ca^2+^]_i_ event was determined relative to the highest detected peak per trace. This served as a criterion to group [Ca^2+^]_i_ oscillations into low-intensity (< 60 % of highest peak) and high-intensity (≥ 60 % of highest peak) events. Per condition or tested stimulus, 4 - 6 independent experiments were performed under identical settings. 30 - 100 individual MIN6 or 60 primary *β*-cells were picked per condition and their responses were averaged. OriginLab, version 8.5 was applied for statistical analysis and plotting.

### Cleaning of plastics in advance of extraction

For the reduction of background signal in MS screens, herein applied 2 mL PCR-grade Eppendorf tubes were cleaned in a MeOH/EtOAc-based procedure. Centrifuge tubes were sonicated in formic acid (10 %, H_2_O) for 20 min, followed by washing with double-distilled H_2_O and with MeOH. The entire procedure was repeated twice, followed by incubation of tubes in MeOH, o/n. On the next day, the same procedure was repeated with EtOAc, instead of MeOH and with incubation of tubes in EtOAc, o/n. After drying, the tubes were applied for sample collections and extractions. Only solvents of the highest analytical grade were applied for extractions and MS analyses.

### Preparation of biological samples for TA extraction

MIN6 cells were grown in 0 60 mm dishes (Nuclon Delta Surface, Thermo Scientific) to form pseudo-islets of 70 - 80% confluence within 5 days after seeding. The growth medium was removed and cells were washed twice with DPBS (1 mL) at RT. MIN6 cells were pre-stimulated with buffer (1.5 mL) for 60 min at 37°C, 5% CO_2_. For priming of MIN6 cells in advance of the application of respective stimuli, identical buffer glucose concentrations were selected in pre-stimulation and stimulation steps. MIN6 cells were incubated in 1.5 mL buffer (+/- stimulus) for 60 min at 37°C, 5% CO_2_.

Following stimulation, MIN6 cells were scraped in 150 µl HEPES buffer, using disposable cell lifters (Thermo Fisher). 50 µl of this were collected for BCA-based protein quantification (Pierce^TM^ BCA Protein assay kit, Thermo Scientific, Rockford, IL, USA). The residual 100 µl were transferred to 2 mL tubes for extraction. 80% MeOH (in H_2_O), supplemented with 2% formic acid was added and samples were incubated for 30 min at -80°C. Samples were concentrated to complete dryness under vacuum at 4°C, o/n. Isotope-labeled TAs were prepared as stock solutions: *β*-PEA (2-phenyl-d_5_-ethylamine, 0.965 mg/mL, EtOH), tryptamine (tryptamine-*α*,*α*,*β*,*β*-d_4_, 6.7 mg/mL, EtOH), synephrine (synephrine-^13^C_2_, ^15^N, 100 μg/mL, DMSO), octopamine (octopamine-d_3_, 1 mg/mL, EtOH) and tyramine (tyramine-d_4_, 1 mg/mL, EtOH), (see Fig. S4). Stocks were combined to generate a heavy internal standard master mix (1 µg/mL in EtOH, in the following: hISDs mix). Heavy-isotope labelled TAs within this master mix were spiked into samples. Stocks of non-labeled TAs were prepared (2 µg/mL in EtOH) which served as references and quality controls in LC-MS measurements.

### MeOH-based TA extraction

80% MeOH in H_2_O (375 µL), acidified with HCl (1% final) and 2 - 4 µL of the hISDs mix (1 µg/mL in EtOH) were added to dried samples (SNs and cell pellets, respectively). Samples were mixed and sonicated for 30 min at 4°C, followed by centrifugation at 4000 x g for 5 min at 4°C. SN and scraped cell samples were combined in pre-cleaned 2 mL tubes. To check for background signal, 2 - 4 µL of the hISDs mix (1 µg/mL in EtOH) were extracted from cleaned 2 mL centrifuge tubes. Also, cleaned tubes without hISDs were included for extractions. Combined organic phases were dried under vacuum at 4°C, o/n. Dry samples were stored under argon at -80°C.

### Liquid chromatography-mass spectrometry (TAs)

Separation of metabolites by liquid chromatography was performed on a Vanquish UHPLC system (Thermo Fisher), equipped with a Luna Omega Polar C18 column (1.6 μm, 100x2.1 mm) at a flow of 0.26 mL/min and at 40 °C. Mobile phase A consisted of 7.5 mM ammonium acetate, supplemented with 0.02 % formic acid (pH 4). Mobile phase B consisted of 7.5 mM ammonium acetate in acetonitrile:H_2_O (95:5). The applied gradient for MeOH/HCl extracted samples is listed in table 2.

**Table 2.** Applied gradient for LC-MS analysis of TA extracts. Mobile phase A consisted of 7.5 mM ammonium acetate, supplemented with 0.02 % formic acid (pH 4), mobile phase B of 7.5 mM ammonium acetate in acetonitrile:H_2_O (95:5).

| Time (min) | % of mobile phase B |
| --- | --- |
| 0 | 5 |
| 1 | 5 |
| 7 | 54 |
| 8 | 90 |
| 11 | 90 |
| 11.5 | 5 |
| 14 | 5 |

Extracted samples were re-suspended in 100 μL mobile phase A. Samples were vortexed (10 s) and centrifuged at maximum speed for 2 min at RT. 100 μL of the clear SN were injected for analysis. MS was performed on a Q-Exactive Plus (Thermo Fisher) high resolution mass spectrometer (HRMS), equipped with an advanced hybrid quadrupole-Orbitrap. Ionization was achieved by electrospray ionization (ESI-MS) in positive mode (spray-voltage: 4 kV). Spectra were acquired as full scans (60 - 900 m/z) with a resolution of 35000 (detailed parameters are listed in table 3). The collected LC-MS data were evaluated using the Xcalibur software tool (Thermo Fisher).

**Table 3.** MS parameters for TA detection.

|  |  |
| --- | --- |
| ESI mode | positive |
| Scan range | full scan (60 - 900 m/z) |
| Resolution | 35,000 |
| Spray voltage | 4 kV |
| S-lens RF level | 65/75 |
| Sheath gas | 30 |
| Auxiliary gas | 5 |
| Probe heater temperature | 300 °C |
| Capillary temperature | 320 °C |
| Injection volume | 20 µL |

## RESULTS

### Trace amines and TAAR1 (ant-)agonists modulate [Ca^2+^]_i_ oscillations and insulin secretion

Our previous work supports the hypothesis of an essential role of autocrine signaling factors for glucose-stimulated *β*-cell activity and insulin secretion, in particular, fatty acids as ligands of GPR40 and extracellular ATP [12,17]. TAAR1 expression was identified in pancreatic *β*-cells, and TAAR1 agonists have been reported to modulate insulin secretion [43]. We therefore suspected ligands of TAAR1 to also participate in *β*-cell crosstalk and autocrine activation.

It is established that the *β*-cell line MIN6 responds well to different levels of external glucose by periodic transients of the cytosolic Ca^2+^ concentration ([Ca^2+^]_i_ oscillations) and insulin secretion, similar to primary *β*-cells [48]. In the present work, we exploited the well-documented correlation between [Ca^2+^]_i_ oscillations and insulin secretion (Supplemental Fig. S1 and [49–51]) as a sensitive readout of *β-*cell response. [Ca^2+^]_i_ oscillations were followed in real-time, using the cell-permeant version of the Ca^2+^-sensitive fluorescent indicator Fluo-4 [52]. Intracellular cAMP levels were monitored as a readout for TAAR1 activity, utilizing an EPAC-based FRET sensor [46]. Pre-washing MIN6 and mouse primary *β-*cells transiently stopped [Ca^2+^]_i_ oscillations (Fig. 1), as described previously [12].

**Figure 1.**
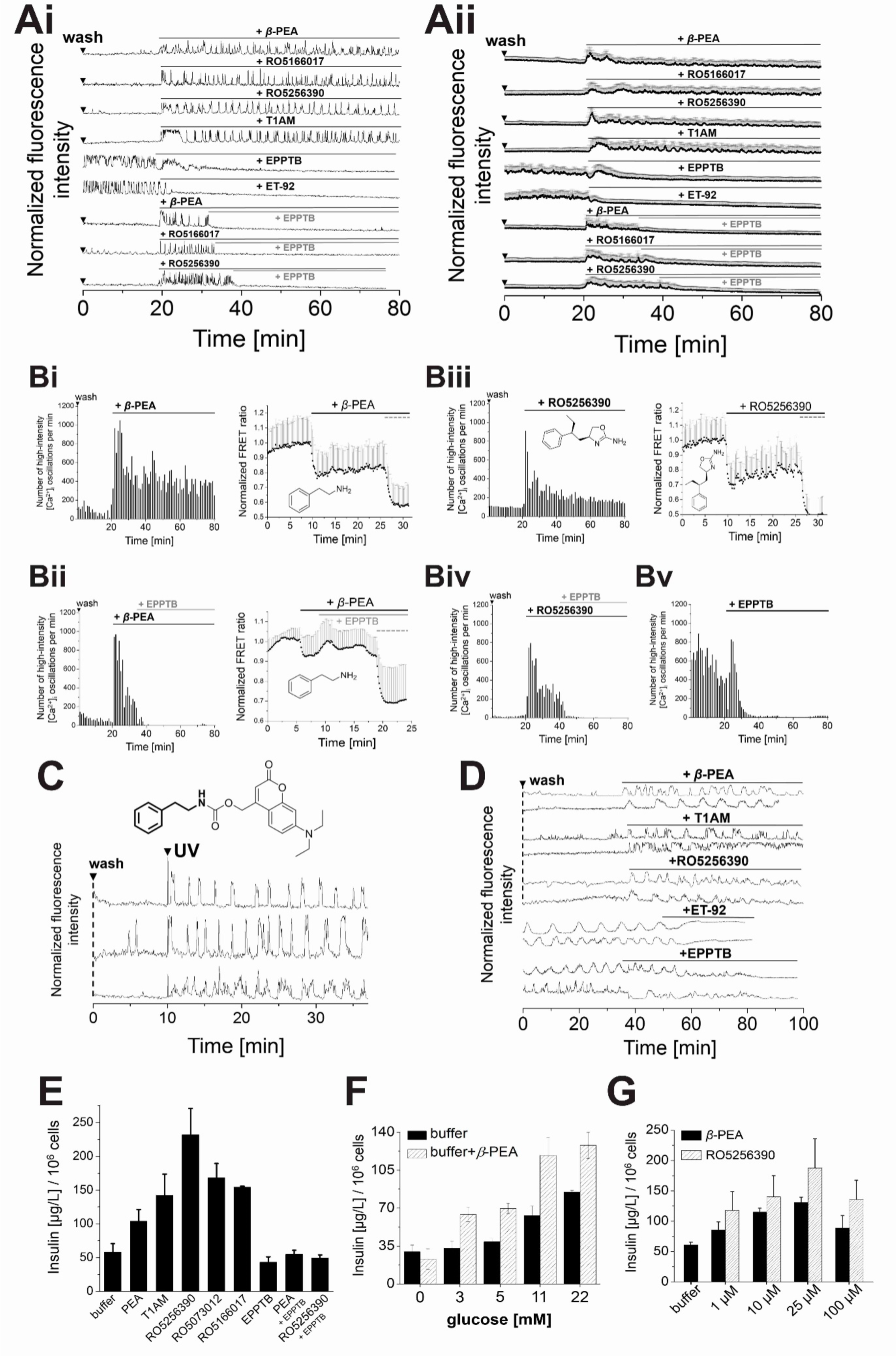
Modulation of [Ca^2+^]_i_ oscillations and insulin secretion by TAAR1 agonists and antagonists in MIN6 and mouse primary *β*-cells. **(A)** Representative single **(Ai)** and averaged **(Aii)** Ca^2+^ traces from MIN6 cells, recorded with the Ca^2+^ indicator Fluo-4. T1AM evoked full stimulation already at 10 µM (11 mM glucose). All other compounds were applied at 25 µM final concentrations in the presence of 11 mM glucose. Suppression of [Ca^2+^]_i_ oscillations and insulin secretion was observed for selective TAAR1 antagonists EPPTB and ET-92 with or without preceding stimulation by TAAR1 agonists. **(B)** Counts of high-intensity [Ca^2+^]_i_ events per 60 s interval (right) along with normalized FRET ratios (acceptor/donor intensity of the EPAC^3rd^ sensor) (left). Binding of cAMP induces induces a robust decrease of the FRET ratio between Venus and mTurquoise fluorescent proteins. A mix of forskolin and IBMX (50 μM, each) was added to cells at the end of each experiment to induce maximum cAMP responses (dashed grey line). Shown are averages of 20 - 30 MIN6 cells (FRET) and 100 MIN6 cells (Ca^2+^ traces). **(C)** Photolysis of caged *β*-PEA by a short UV pulse (λ = 375 nm) instantaneously started [Ca^2+^]_i_ oscillations in pre-washed MIN6 cells. **(D)** Representative Ca^2+^ traces from mouse primary *β*-cells. Compounds were applied at 25 µM final concentrations in the presence of 5 mM glucose. Agonistic effects on insulin secretion were reversed by the antagonists ET-92 and EPPTB. **(E)** Levels of secreted insulin from 10^6^ MIN6 cells as determined by ELISA. *β*-PEA, RO5166017, RO5256390 and the endogenous agonist T1AM stimulated insulin secretion 3- to 5-fold compared to buffer levels. **(F)** Stimulation of insulin secretion by *β*-PEA (25 μM) in the presence of different glucose concentrations, as indicated. **(G)** Concentration-dependent stimulation of GSIS from MIN6 cells by *β*-PEA or RO5256390 with applied concentrations, as indicated. Washes are indicated by ▾. If not indicated otherwise, experiments were performed in the presence of 11 mM glucose. Insulin experiments were performed in quadruplicate (*P<0.05 and **P<0.01. All unmarked events in E, F and G are not statistically significant = P>0.05. ANOVA, with repeated measures as necessary). Ca^2+^ traces were recorded from cells that were stained with the Ca^2+^ indicator Fluo-4. Shown are averages of 100 MIN6 cells.

The *β*-PEA skeleton is an essential molecular feature that all TAAR1 agonists share. This structural element is also present in the decarboxylated skeleton of thyroid hormone molecular derivatives, known as thyronamines, such as T1AM [53]. The addition of TAAR1 agonists *β*-PEA or T1AM to pre-washed MIN6 or mouse primary *β*-cells immediately recovered [Ca^2+^]_i_ oscillations (Fig. 1B) and increased intracellular cAMP levels (Fig. 1B).

As TAAR1 has been described to play a significant role in the modulation of psychiatric disorders, such as schizophrenia, Parkinsońs disease, or drug addiction [32], several full (RO5166017, RO5256390 [29,30]) or partial (RO5073012 [38]) TAAR1 agonists have been designed and made commercially available. Treatment of pre-washed MIN6 cells with TAAR1 agonists RO5166017, RO5256390, and the partial agonist RO5073012 (all at 25 µM) immediately induced continuous [Ca^2+^]_i_ oscillations and cAMP formation (Fig. 1A+B), indicating that the compounds replaced trace amines that were removed by the wash procedure.

#### Spatio-temporal control over TAAR1 activation

To trigger TAAR1 activation with high spatio-temporal control, we synthesized a photo-caged *β*-PEA (Fig 1 C). For this, the photoactivatable “caging” group 7-diethylamino-4-methylene-coumarin was attached to the primary amine of *β*-PEA via carbamate formation (see supplemental data for details on the synthesis). The photo-labile coumarin cage was intended to mask *β*-PEA and to prevent receptor interaction and oxidation by monoamine oxidases (MAO). MIN6 cells were pre-incubated with caged *β*-PEA and washed. Application of a short UV-pulse (375 nm) immediately induced continuous [Ca^2+^]_i_ oscillations (Fig 1 C), indicating that cellular activity can be modulated by the stimulation of TAAR1.

Stimulation of TAAR1 increased insulin secretion 3- to 5-fold, with the most potent stimulation by artificial ‘RO-agonists’, followed by the endogenous TAAR1 agonists T1AM and *β*-PEA (Fig 1 E). Stimulation of MIN6 cells by *β*-PEA (25 µM) yielded significantly higher levels of insulin at stimulatory glucose concentrations, compared to sub-stimulatory glucose levels (Fig. 1 F). To check whether TAAR1-mediated potentiation of GSIS was concentration-dependent, MIN6 cells were stimulated with increasing concentrations of *β*-PEA and RO5256390 in the presence of 11 mM glucose. Insulin secretion reached maximum levels in the presence of 25 µM *β*-PEA (Fig. 1 G).

TAAR1-antagonists EPPTB [54] and ET-92 [39] significantly reduced spontaneous [Ca^2+^]_i_ oscillations and insulin secretion from MIN6 and mouse primary *β*-cells (Fig 1 A+B, E). This indicated basal (constitutive) TAAR1 activation by endogenous ligands and confirmed EPPTB and ET-92 as antagonists. Similarly, after TAAR1 stimulation we observed comparable effects of antagonists on [Ca^2+^]_i_ oscillations and insulin secretion for MIN6 cells (Fig. 1 A+B).

The reversal of agonist-mediated effects by the selective antagonist EPPTB has been defined as a criterion to identify effects that are mediated by TAAR1 stimulation [54]. *β*-PEA and RO5256390-mediated effects on insulin secretion were overruled down to buffer levels in the presence of EPPTB (Fig. 1 E).

### TAs and their derivatives show differences in their stimulatory potencies

By comparing the stimulatory potencies of different TAs on [Ca^2+^]_i_ oscillations in MIN6 cells, we found that *β*-PEA and tyramine were more potent than synephrine and octopamine (Supplemental Fig. S2 A+Bi-v). Based on their effects on [Ca^2+^]_i_ responses, we determined a rank order of potency: *β*-PEA > *p*-tyramine > tryptamine >> *p*-synephrine ≈ *p*-octopamine (Supplemental Fig. S2A+Bi-v), in line with literature reports [55,56]. Corresponding amino acids did not stimulate [Ca^2+^]_i_ oscillations in pre-washed MIN6 cells (Supplemental Fig. S2 A+B vi-viii).

Also, we observed that TAAR1 exhibited broad ligand specificity, as [Ca^2+^]_i_ oscillations and insulin secretion were not only stimulated by amines defined as TAs, but also by an array of other primary amines. Therefore, we profiled a selection of compounds structurally related to *β*-PEA-for their potency to stimulate TAAR1, e.g. with specific modifications of the aromatic moiety and ethylene chain of *β*-PEA (Supplemental Fig. S3). We found that the potency to evoke high-intensity [Ca^2+^]_i_ oscillations decreased with increasing complexity of the amine. Secondary amines, such as *N*-methyl-PEA and synephrine, were less potent than the corresponding primary amines (Supplemental Fig. S3).

### Inhibition of TA biochemical pathways modulates endogenous TA levels, [Ca^2+^]_i_ oscillations and insulin secretion

Biosynthetic pathways for TAs have been described for neuronal tissue, where *β*-PEA, tyramine and tryptamine are generated by decarboxylation from precursor amino acids. This rate-limiting step is performed by AADC and followed by additional biosynthetic steps for the generation of secondary amines synephrine, *N-*methyltyramine and *N*-methylphenylethylamine or octopamine [57]. TAs are rapidly inactivated in a single enzymatic oxidation step by monoamine oxidase (MAO). While TAs are metabolized by both MAO-A/B isoforms, *β-*PEA is primarily converted by MAO-B [58–61]. MAOs show high turnover rates, leaving TAs with half-lives in the range of ∼30 s [60]. As MAO-B also metabolizes dopamine, inhibitors of MAOs are applied for the treatment of depression and Parkinson’s disease [62].

We turned to mass spectrometry (MS) to identify and quantify levels of endogenous TAs from MIN6 cells. For this, the supernatant of MIN6 cells was collected, dried and combined with scraped MIN6 cells for MeOH / HCl extraction in the presence of stable heavy isotope-labeled internal standards (hISDs, Supplemental Fig. S4). MS analysis of MIN6 cell supernatants identified a variety of primary amines including *β*-PEA and tyramine in trace amounts (Fig. 2A).

**Figure 2.**
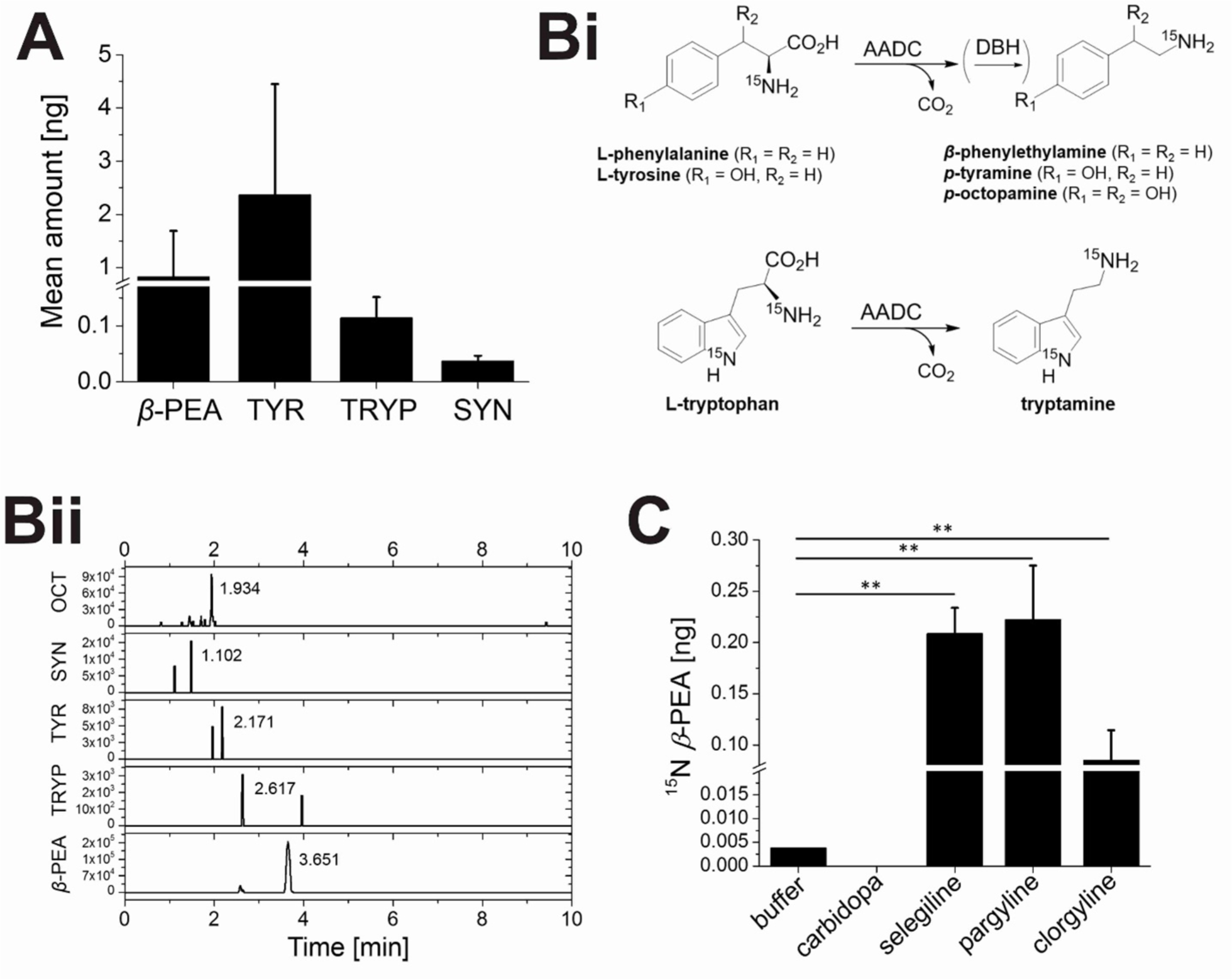
TAs are biosynthesized in MIN6 cells and detectable levels are modulated by exogenous stimuli. MIN6 cells were grown to 70-80 % confluency and treated with exogenous stimuli reported to modulate insulin secretion by increasing glucose concentration (0-22 mM glucose) and applying high potassium levels (30 mM KCl). TAs were extracted from stimulated cells and analyzed by LC-MS performed with a Vanquish UHPLC system (Thermo Fisher), followed by mass-based separation with a Q-Exactive Plus (Thermo Fisher) high resolution mass spectrometer (HRMS) equipped with an advanced hybrid quadrupole-Orbitrap. If not stated otherwise, buffer refers to buffer supplemented with 11 mM glucose, which served as the control condition for most experiments. **A:** mean TA concentration [ng] detected in MIN6 cells stimulated with 11 mM glucose; PEA (n = 3), TYR (n = 4), TRYP (n = 2) and SYN (n = 2), with n being the number of replicates for each TA. **Bi:** MIN6 cells grown in the presence of ^15^N-labeled L-tyrosine, L-phenylalanine and L-tryptophan were found to synthesize and secrete ^15^N-labeled TAs based on the depicted schematic biosynthetic pathway for TAs. **Bii:** exemplary intensity-over-time peaks for all TAs as detected in control samples; the number represents the respective retention time. **C:** MIN6 cells treated with inhibitors for key metabolic enzymes involved in both TA biosynthesis and degradation (carbidopa = AADC inhibitor, seleglinide = pargyline = MAO-B inhibitors and clorgyline = MAO-A inhibitor), were found to adjust detectable TA levels based on the applied condition, exemplary shown for ^15^N-PEA. All experiments were performed in quadruplicate, error bars represent ±SEM. *p<0.05, **p<0.01 and ***p<0.001, all unmarked events were not statistically significant.

Sub-culturing MIN6 cells in the presence of heavy-labeled canonical amino acids, such as ^15^N-phenylalanine, –tyrosine, and -tryptophan lead to the identification of heavy-labeled primary amines in cellular supernatants, which are most likely produced by AADC-mediated decarboxylation (Fig. 2B).

To understand biosynthetic pathways of TAs and the role of endogenous TAs for cellular activity, we inhibited key enzymes of TA biosynthesis and degradation pathways in MIN6 cells cultured in the presence of ^15^N-phenylalanine to generate ^15^N-*β*-PEA. MS analysis showed that ^15^N-*β*-PEA levels were significantly increased in MAO inhibitor-treated MIN6 cells (Fig. 2 C). Inhibition of AADC by carbidopa reduced ^15^N-*β*-PEA levels below the detection limit (Fig. 2 C). Therefore, MIN6 cells are capable of synthesizing and degrading *β*-PEA, and likely several other TAs, suggesting TAs as potential autocrine signaling molecules.

To further investigate the potential of Tas as autocrine signals, MIN6 cells were treated with recombinant MAO-B. The immediate reduction in [Ca^2+^]_i_ oscillations (Fig. 3A+Bi) following MAO-B addition supported the hypothesis of an essential role of primary amines for fully functional autocrine signaling in *β*-cells and suggests a rapid exchange between intra- and extracellular TAs.

**Figure 3.**
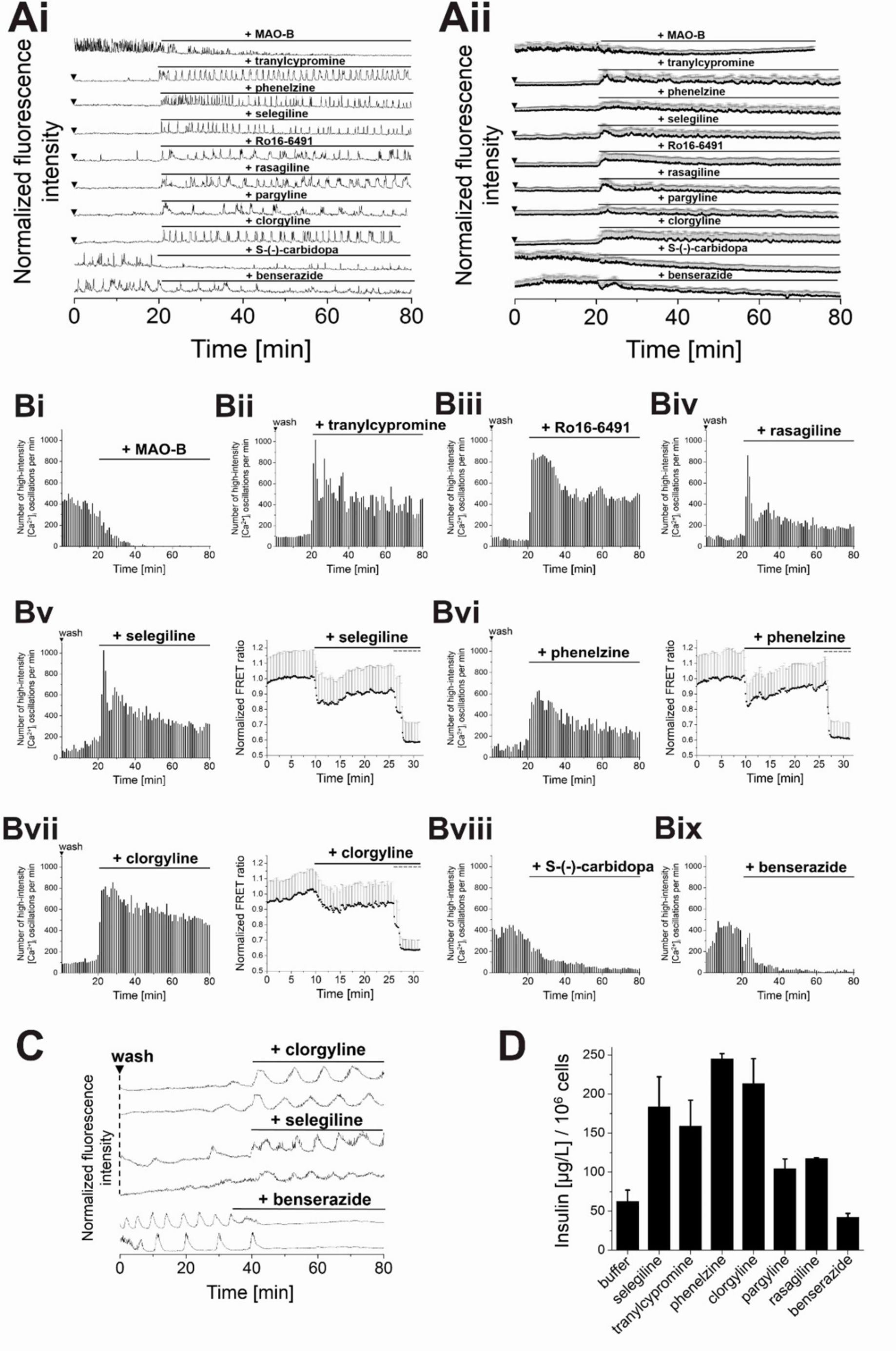
Inhibition of TA biochemical pathways modulates intracellular [Ca^2+^]_i_ oscillations, cAMP levels and insulin secretion. **(A)** Representative single **(Ai)** and averaged **(Aii)** Ca^2+^ traces from MIN6 cells, recorded with the Ca^2+^ indicator Fluo-4. **(B)** Counts of high-intensity [Ca^2+^]_i_ events per 60 s interval. Intracellular cAMP levels were monitored via the FRET sensor EPAC 3^rd^ (mTurquoise-Epac(CD,delDEP)-^cp173^Venus-Venus. **(A, Bi)** [Ca^2+^]_i_ oscillations in glucose stimulated MIN6 cells gradually decreased in the presence of recombinant MAO-B (0.3 U). **(A, Bii-vii)** Inhibition of MAO by selective MAO-A/B inhibitors potentiated intracellular [Ca^2+^]_i_ oscillations and cAMP formation in pre-washed MIN6 cells and **(C)** primary mouse *β*-cells. Also, insulin secretion increased in the presence of MAO-inhibitors. **(Bviii+ix)** Inhibition of AADC reduced [Ca^2+^]_i_ oscillations in glucose-stimulated MIN6 cells and in **(C)** mouse primary *β*-cells. **(D)** Insulin secretion from MIN6 cells was significantly decreased by MAO inhibition. MAO-inhibitors were applied at 25 μM, AADC inhibitors at 50 μM in the presence of 11 mM glucose. Shown are averages of 100 MIN6 cells. Insulin experiments were performed in quadruplicate (*P<0.05 and **P<0.01. All unmarked events in D are not statistically significant = P>0.05. ANOVA, with repeated measures as necessary). Error bars present SD. Washes are indicated by ▾. Imaging and insulin experiments were performed in the presence of 11 mM glucose.

TA biosynthetic inhibitors were applied to glucose-stimulated MIN6 and mouse primary *β*-cells. The effects were followed by monitoring [Ca^2+^]_i_ oscillations, intracellular cAMP levels, and insulin secretion. The number of high-intensity [Ca^2+^]_i_ oscillations and the levels of cAMP and insulin secretion were significantly increased in the presence of MAO inhibitors, regardless of their mechanism of action or specificity towards MAO-A or -B (Fig. 3A+B). As the modulation of biochemical pathways for the synthesis and degradation of TAs directly translated into changes in *β*-cell activity and insulin secretion, we inferred high metabolic turnover rates of TAs and autocrine feedback. Similarly, pre-washed mouse primary *β*-cells responded to the inhibition of MAO with increased [Ca^2+^]_i_ oscillations, while inhibition of AADC by benserazide decreased [Ca^2+^]_i_ oscillations (Fig. 3 C).

### Psychotropic drugs stimulate [Ca^2+^]_i_ oscillations and insulin secretion via TAAR1

Addictive drugs, such as (meth-)amphetamine and 3,4-methylenedioxymethamphetamine (MDMA) share the same scaffold with *β-*PEA and have been described to act as potent TAAR1 agonists in the CNS [25]. Therefore, we tested a spectrum of psychotropic drugs for their effects on [Ca^2+^]_i_ oscillations and insulin secretion from MIN6 and mouse primary *β*-cells (Fig. 4). Compounds with agonistic effects included antidepressants dibenzepine, idazoxan, and nomifensine, the benzodiazepine flurazepam, the antiparkinson agent diphenylhydramine, as well as ergoline derivatives lisuride (antiparkinson agent) and methysergide (migraine treatment) (Fig. 4). Hence, we also expected amphetamine-mediated stimulation of *β*-cells via TAAR1. Administration of amphetamine to pre-washed MIN6 cells immediately induced [Ca^2+^]_i_ oscillations and increased insulin levels (Fig 4A-C). Drug-induced [Ca^2+^]_i_ oscillations were suppressed by the addition of the TAAR1 antagonist EPPTB, confirming that amphetamines exert their effects predominantly through TAAR1 (Fig. 4). Our observations are in line with literature reports, according to which administration of amphetamine produces a rapid increase of immune assay detectable plasma insulin in mice and rats [63]. Early reports hypothesized that insulin is released by a neural mechanism within the pancreas, but a *β*-cell-derived mechanism was not ruled out [63]. Based on our results, we suggest a direct impact of these psychoactive compounds on *β*-cell physiology, independent from CNS effects.

**Figure 4.**
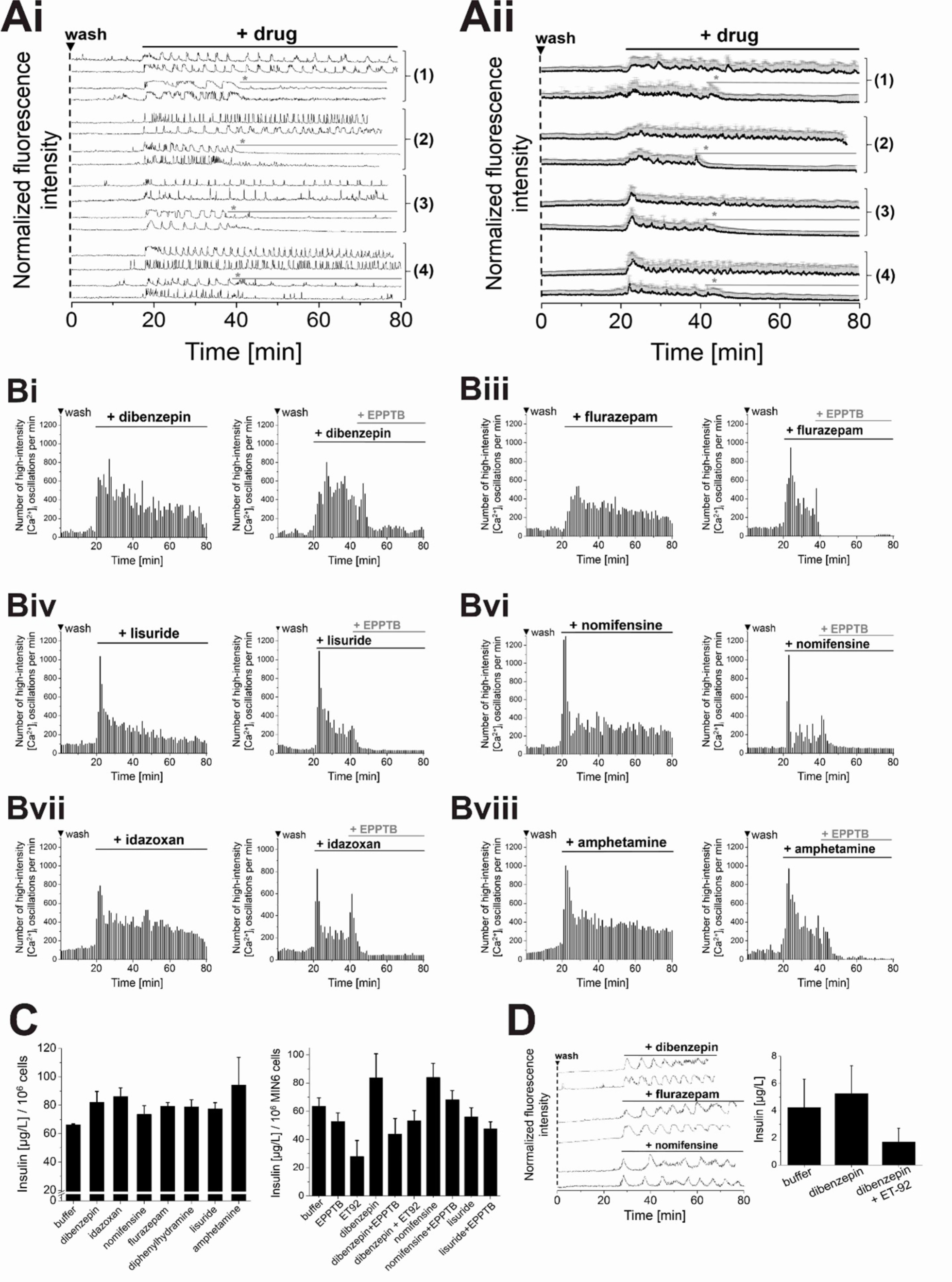
Psychotropic drugs modulate [Ca^2+^]_i_ oscillations in MIN6 and primary mouse *β*-cells via TAAR1. Representative single **(Ai)** and averaged **(Aii)** Ca^2+^ traces from MIN6 cells, recorded with the Ca^2+^ indicator Fluo-4. **(Bi – Bvii)** Counts of detected high-intensity [Ca^2+^]_i_ events per 60 s interval. Differential stimulation of [Ca^2+^]_i_ oscillations in pre-washed MIN6 cells was observed upon addition of antidepressants **(Ai+ii (1)**, **Bi)** dibenzepin, **(Bii)** idazoxan and **(Biii)** nomifensine, the benzodiazepine **(Ai+ii (2)**, **Biv)** flurazepam, the antiparkinsonian agents **(Ai+ii (3)**, **Bv)** lisuride and **(Bvi)** diphenylhydramine, as well as **(Ai+ii (4)**, **Bvii)** amphetamine. Intracellular cAMP levels following amphetamine treatment were monitored via the FRET sensor EPAC 3^rd^ (mTurquoise-Epac(CD,delDEP)-^cp173^Venus-Venus. Drug-induced stimulation of [Ca^2+^]_i_ oscillations was reduced by the addition of the TAAR1 antagonist EPPTB (50 µM) confirming TAAR1 action. **(C)** Insulin secretion from MIN6 cells in the presence of psychoactive drug and in combination with EPPTB. **(D)** [Ca^2+^]_i_ traces and insulin secretion from primary mouse *β*-cells after stimulation with psychoactive drugs, as indicated. Compounds were applied at 25 µM in the presence of 11 mM glucose. Shown are averages of 100 MIN6 cells. Insulin experiments were performed in quadruplicate. Insulin experiments were performed in quadruplicate (*P<0.05. All unmarked events in C and D are not statistically significant = P>0.05. ANOVA, with repeated measures as necessary). Error bars present SD. Washes are indicated by ▾.

## DISCUSSION

The detection of TAAR1 mRNA in peripheral tissue, in particular in mouse and human pancreatic *β*-cells as well as in the cell lines INS-1 and MIN6 initially indicated that TAs might play a role in the periphery [40,42]. However, TA levels in plasma are known to be in the sub-nanomolar range [64,65]. Therefore, it was unclear whether their concentration would be sufficient to activate peripheral TAAR1.

Our initial experiments showed that TAs modulated calcium signaling and insulin secretion in *β*-cells over a broad concentration range. This starting point led us to the hypotheses that 1) TAs are produced by *β*-cells in an autocrine fashion and 2) local TA levels in the islet reach high levels and are quite transient. Therefore, this signaling system might act quasi-autonomous similar to the recently described fatty acid- and ATP-based autocrine regulation systems in the islets of Langerhans [12,17]. This hormone-GPCR signaling system differs significantly from more thoroughly described organism-wide adrenergic and muscarinic signaling. The relevance of the TA/TAAR1 system is highlighted by a recent report that single-nucleotide variants in TAAR1 were connected to disturbed glucose homeostasis in overweight/obese patients [65].

Following our hypothesis, we provide evidence suggesting that MIN6 cells possess the necessary metabolic machinery to transform labeled precursor amino acids into labeled TAs, as indicated by the identification of ^15^N-labeled TAs in cell supernatants by LC-MS. In addition, the immediate effect of the antagonists EPPTB and ET-92 shows that there is a persisting TA tone that is essential for productive signaling and secretion. Considering the restricted extracellular space available within an islet, concentrations of secreted molecules can build up rather quickly, easily reaching concentrations in the µM range [14]. Similarly, Revel *et al.* reported TAAR1 activation potencies of RO5256390 and RO5166017 that were in comparable ranges to those of *β*-PEA [29,30], based on cAMP measurements. The micromolar concentrations of antagonists (EPPTB and ET-92) needed to block the TA effect further support this level of accumulation. The extra- and potentially -intracellular local TA concentration seems to be limited by MAO activity and potentially by amine uptake transporters such as the dopamine transporter (Fig. 5).

**Figure 5.**
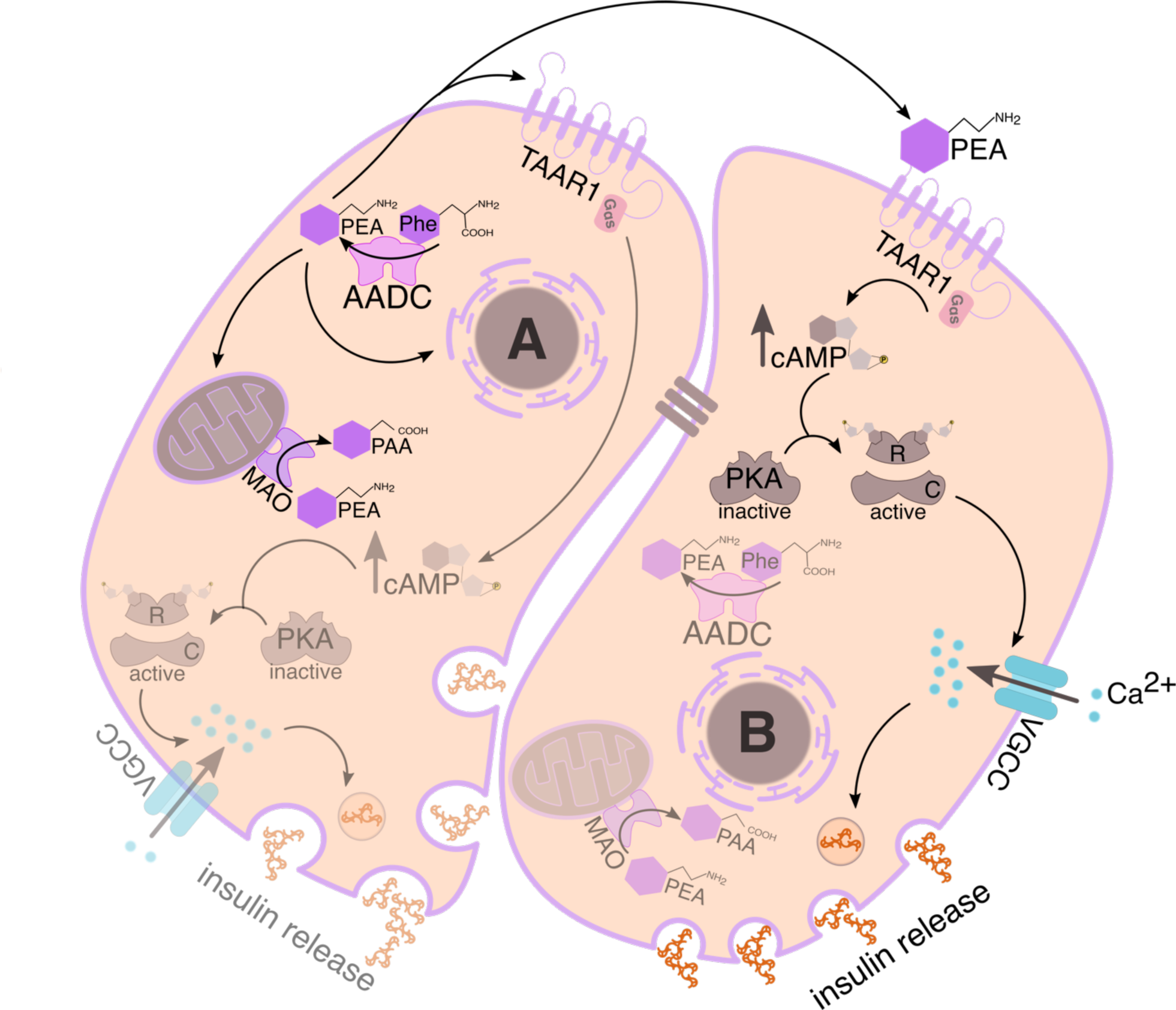
Proposed mechanism of TAAR1 signaling in pancreatic *β*-cells. TAAR1 is known to reside on endomembranes and likely the plasma membrane. It is stimulated by endogenous trace amines (TAs) such as PEA in an autocrine (*β*-cell A) or paracrine (*β*-cell B) manner. Cell A highlights the biosynthetic machinery for the synthesis and degradation of TAs within *β*-cells. Aromatic amino acid decarboxylase (AADC) catalyzes the conversion of aromatic amino acids to TAs (exemplary shown is the conversion of phenylalanine (Phe) to PEA). PEA might activate TAAR1 of the same cell (autocrine) or on neighboring cells (paracrine). TAs are quickly degraded by monoamine oxidase (MAO), converting PEA into phenylacetic acid (PAA). Cell B highlights the signaling cascade initiated once TAs have activated TAAR1 and subsequently G_αs_, leading to the generation of cAMP which binds to and activates PKA (grey, regulatory subunit R). The liberated catalytic (C) subunit phosphorylates voltage-gated Ca^2+^ channels (VGCC, blue) and hence increases the influx of Ca^2+^ ions (cyan). Increased levels of [Ca^2+^]_i_ trigger the fusion of insulin granules with the PM and insulin secretion. For simplicity, glucose as the main stimulus for insulin release as well as the ER as an internal Ca^2+^ source have been omitted. *β*-cells are electrically coupled by gap junctions (brown bars).

Interestingly, some MAO inhibitors that are prescribed for the treatment of depression or Parkinsońs disease have been reported to come with side effects, namely significant weight gain [67]. Other antidepressants including tricyclic drugs are well known for the same side effects. While these might be due to effects on the central nervous system, we demonstrate here that most drugs that feature an aromatic moiety at a distance of two or three carbons from a protonatable amine, similar to PEA, show comparable effects in calcium signaling and insulin secretion. The only exceptions were molecules with heavily convoluted structures such as ergometrine, dihydroergotamine, D-lysergic acid diethylamide, and bromocriptine.

The combined results make TAAR1 an attractive therapeutic target for the pharmacological modulation of *β-*cell activity and insulin secretion. The existing antagonists, EPPTB and ET-92 might serve as a good starting point as they significantly limit signaling and secretion in *β*-cells, although *in vivo* application of EPPTB might be limited by its high clearance [32,68].

As TAs might shuffle in and out of cells passively or by secretion and transporters, it remains unclear where the receptors are located and active. The application of caged PEA that accumulated in the cells’ endomembranes supported the idea that intracellular signaling via TAAR1 is relevant, maybe even dominant. However, further experimentation is required to answer this question conclusively. Similarly, it is unclear if the positively charged TAs are released via secretory granules or can leave cells as soon as cells are depolarized.

## CONCLUSION

Our finding that TAs are essential regulators for calcium signaling and insulin secretion in *β*-cells adds another autocrine factor to the suite of autocrine signaling molecules in the islets of Langerhans. Future research will show how extracellular signaling molecules such as ATP and fatty acids will substitute or coordinate the function of TAs in *β*-cell signaling and how the extracellular factors will affect signaling in the other cell types within the islet. The strong effects of many classical drugs with aromatic amine moieties on TAAR1 and *β*-cell signaling need to be accommodated in future pharmacological studies. In addition, TAAR1 itself might evolve as an important drug target to treat metabolic diseases such as diabetes in the periphery in the future. To evaluate these possibilities, new agonists need to be developed that are unable to pass the blood-brain barrier and hence lack effects on the CNS.

## ACKNOWLEDGEMENT

This research was supported by Transregios 83 and 186, funded by the Deutsche Forschungsgemeinschaft. D.A.Y. acknowledges the IOCB installation grant and C.S. funding from EMBL and OHSU. No potential conflicts of interest to this article were reported.

## Author Contributions

S. H. designed the research; performed experiments; analyzed the data; and wrote, reviewed, and edited the manuscript. K. K. performed experiments; analyzed the data; and reviewed and edited the manuscript. A. L. synthesized caged PEA. M. C. and J. R. performed experiments and analyzed data. D. G. introduced us to TAAR1 and provided samples. D.A.Y. and C.S. designed the research and wrote, reviewed, and edited the manuscript. C.S. is the guarantor of this work and, as such, had full access to all the data in the study and takes responsibility for the integrity of the data and the accuracy of the data analysis.

