## Supplement Data for "Trace Amines are Essential Metabolites for the Autocrine Regulation of *β*-Cell Signaling and Insulin Secretion"

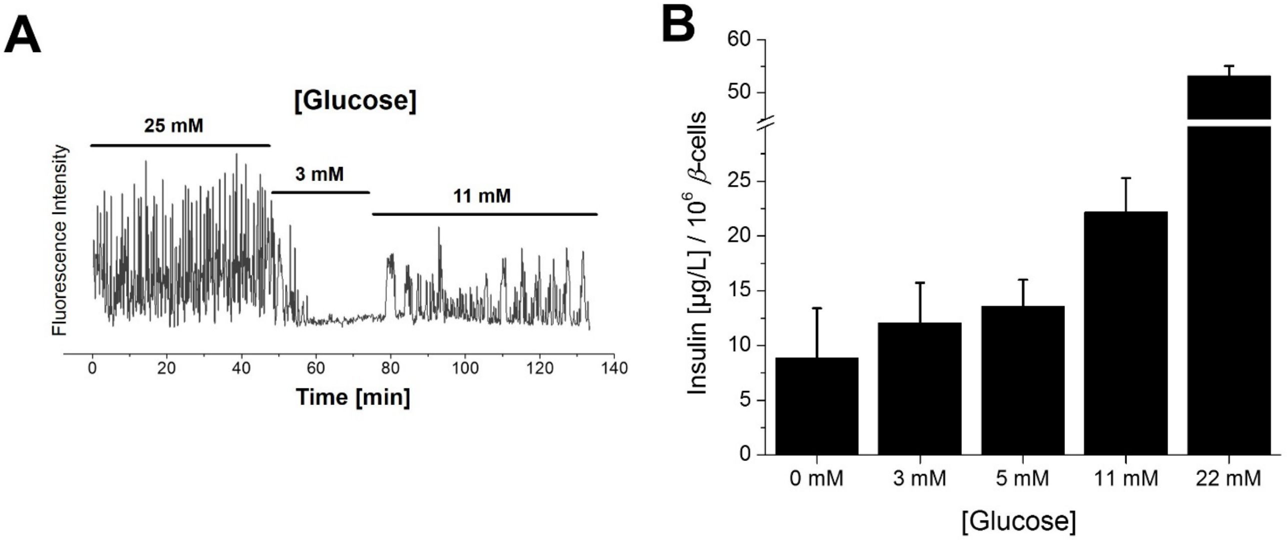


**Figure S1. Highly oscillatory $Ca^{2+}$ responses to various levels of extracellular glucose.** Dependence of $[Ca^{2+}]_i$ oscillations in MIN6 cells and insulin secretion on the applied buffer glucose concentration. **(A)** In order to test the dynamics of the $[Ca^{2+}]_i$ response, MIN6 cells were treated with 3 mM, 11 mM and 25 mM glucose in buffer. No, intermediate or strong $[Ca^{2+}]_i$ oscillations were observed, respectively. For $[Ca^{2+}]_i$ imaging, the $Ca^{2+}$ sensor R-GECO was transiently overexpressed in MIN6 cells. **(B)** Insulin levels at sub-stimulatory glucose concentrations (0 – 5 mM) were significantly lower compared to those observed at stimulatory glucose concentrations (11 and 22 mM). Experiments for the determination of insulin secretion were performed in quadruplicates.

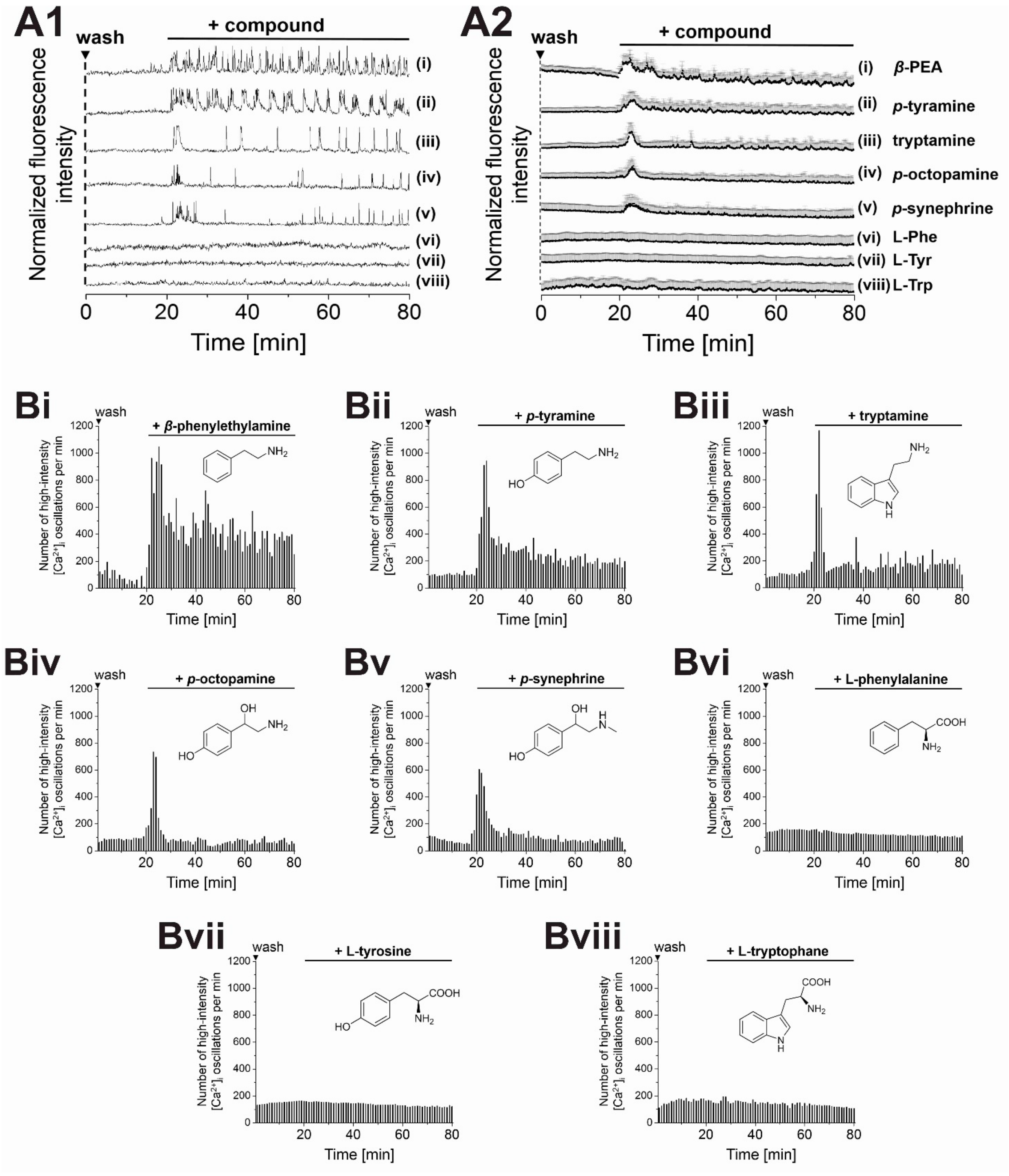


**Figure S2. TAs stimulate $[Ca^{2+}]_i$ oscillations in pre-washed MIN6 cells.** Representative single **(A1)** and averaged **(A2)** $Ca^{2+}$ traces from MIN6 cells, recorded with the $Ca^{2+}$ indicator Fluo-4. **(B)** Number of high-intensity $[Ca^{2+}]_i$ events per 60 s interval. Effects of amines on $[Ca^{2+}]_i$ oscillations decreased with increasing modifications on the aliphatic side chain or amino group. **(A+B i-v)** Based on $[Ca^{2+}]_i$-responses, a rank order of potency was determined for TAs: *β*-PEA > *p*-tyramine > tryptamine >> *p*-synephrine ≈ *p*-octopamine. **(A+B vi-viii)** Corresponding amino acids served as controls. Compounds were applied at 25 µM final concentration in the presence of 11 mM glucose. Shown are averaged traces from 100 MIN6 cells.

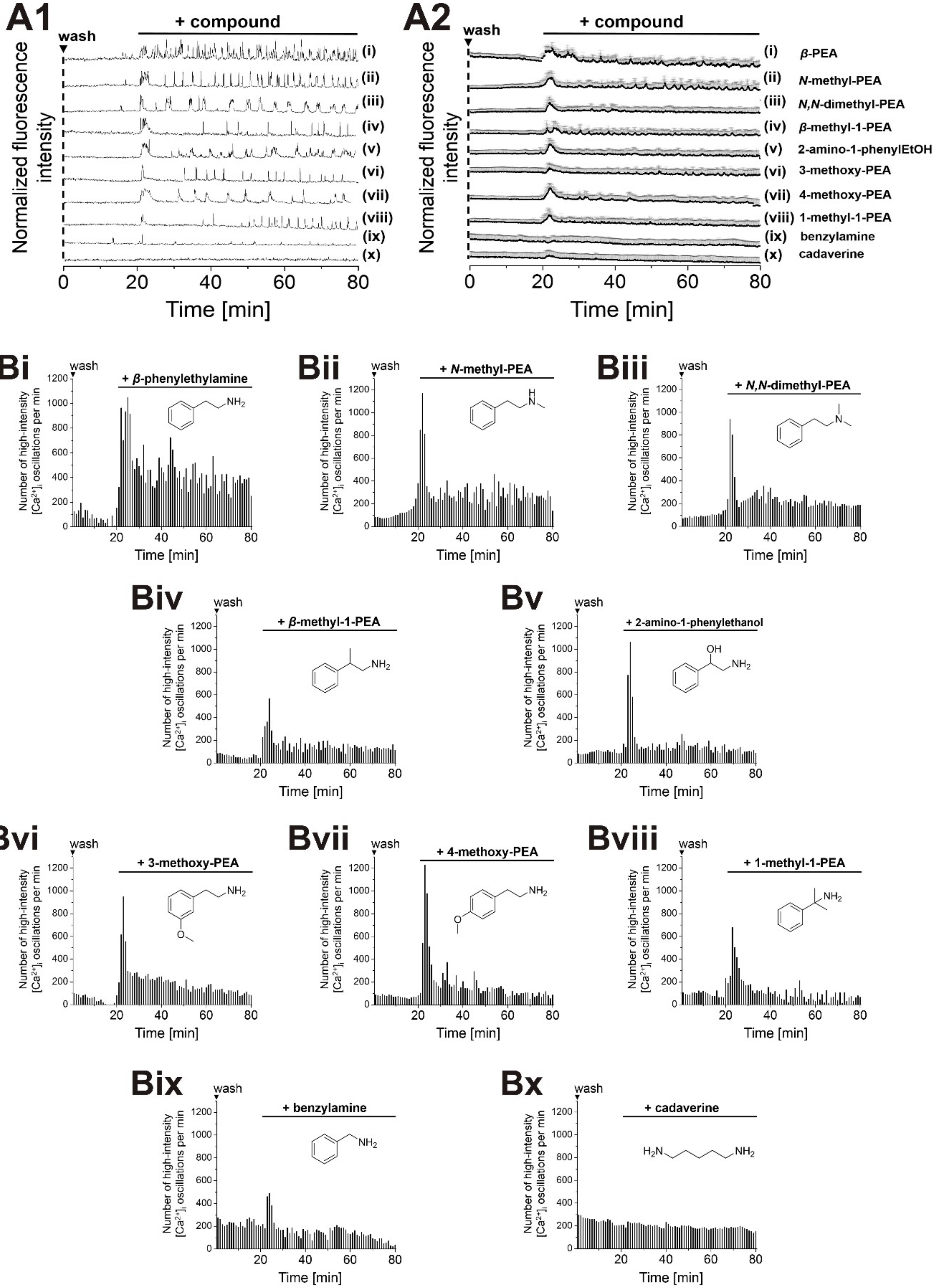
A1
A2
wash
+ compound
Normalized fluorescence intensity
Time [min]
(i) β-PEA
(ii) N-methyl-PEA
(iii) N,N-dimethyl-PEA
(iv) β-methyl-1-PEA
(v) 2-amino-1-phenylEtOH
(vi) 3-methoxy-PEA
(vii) 4-methoxy-PEA
(viii) 1-methyl-1-PEA
(ix) benzylamine
(x) cadaverine
Bi
+ β-phenylethylamine
Bii
+ N-methyl-PEA
Biii
+ N,N-dimethyl-PEA
Biv
+ β-methyl-1-PEA
Bv
+ 2-amino-1-phenylethanol
Bvi
+ 3-methoxy-PEA
Bvii
+ 4-methoxy-PEA
Bviii
+ 1-methyl-1-PEA
Bix
+ benzylamine
Bx
+ cadaverine
Number of high-intensity [Ca2+]i oscillations per min
Time [min]

**Figure S3. Differential stimulation of $[Ca^{2+}]_i$ oscillations in MIN6 cells by aromatic amines.** Representative single **(A1)** and averaged **(A2)** $Ca^{2+}$ traces from MIN6 cells, recorded with the $Ca^{2+}$ indicator Fluo-4. **(B)** Number of high-intensity $[Ca^{2+}]_i$ events per 60 s interval. A selection of (partly *β*-PEA-related) amines was profiled for their potency to stimulate $[Ca^{2+}]_i$ oscillations. Compounds were applied at 25 µM final concentration in the presence of 11 mM glucose. Shown are averaged traces from 100 MIN6 cells.

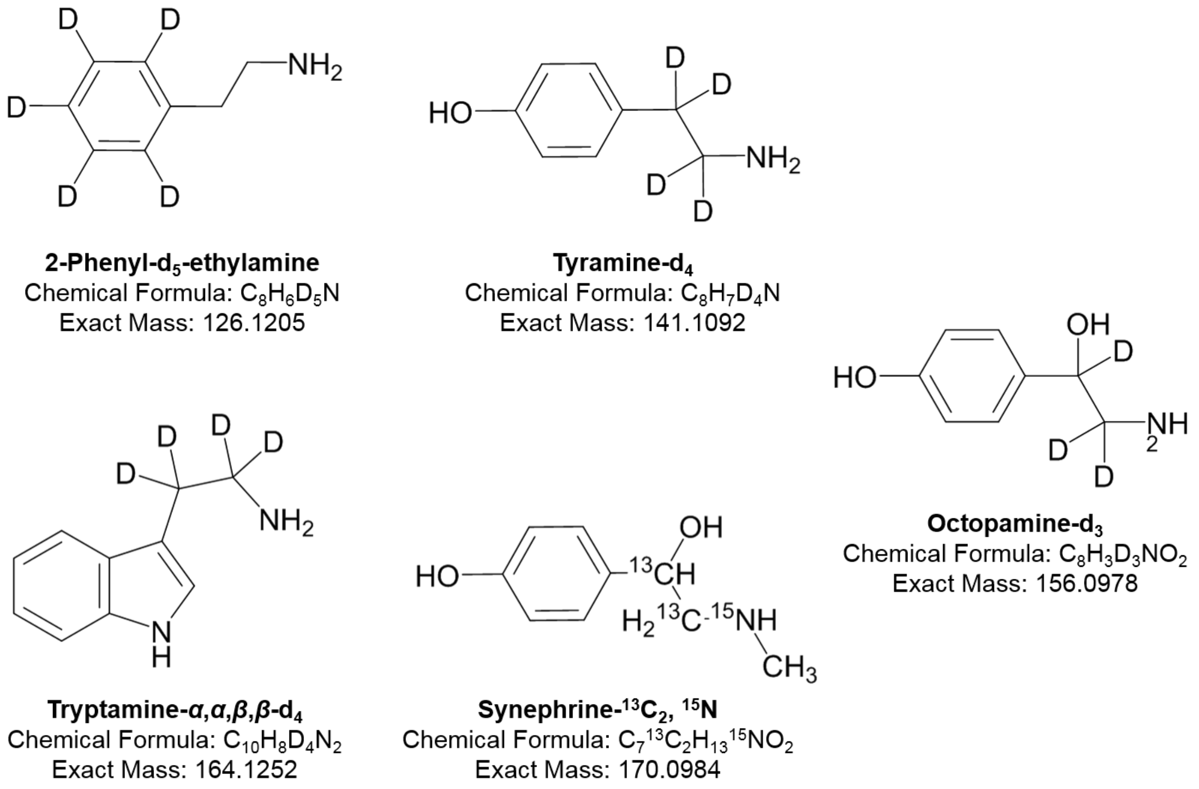


**Figure S4. Overview of stable heavy isotope-labelled TAs that were applied as internal standards for the quantification of TA levels in MS screens.**

**Chemical synthesis of caged phenylethylamine (cg-PEA).**

All chemicals were obtained from commercial sources (Acros, Sigma Aldrich, or Enzo) and were used without further purification. Solvents for chromatography (HPLC grade) were obtained from VWR and dry solvents were obtained from Sigma. Deuterated solvents were obtained from Deutero GmbH, Karlsruhe, Germany. All reactions were carried out using dry solvents under an inert atmosphere unless otherwise stated in the respective experimental procedure. TLC was performed on pre-coated plates of silica gel (Merck, 60 F254) using UV light (254 or 366 nm) or a solution of $KMnO_4$ in $H_2O$ (1.5 g $KMnO_4$, 10 g $K_2CO_3$ and 1.25 mL 10% NaOH in 200 mL of $H_2O$) for analysis. HPLC was performed on a 1260 Infinity system from Agilent Technologies equipped with an EC 250/4 NUCLEODUR 100-5 C18ec analytical column. $^{1}H$- and $^{13}C$-NMR spectra were measured on a 400 MHz Bruker UltraShield$^{TM}$ spectrometer. Chemical shifts of $^{1}H$- and $^{13}C$-NMR-spectra are referenced indirectly to tetramethylsilane. Coupling constants (*J* values) are given in Hz and chemical shifts in ppm. Splitting patterns are designated as follows: s, singlet; d, doublet; t, triplet; q, quartet; m, multiplet; b, broad. $^{13}C$-NMR-spectra were broadband hydrogen decoupled.

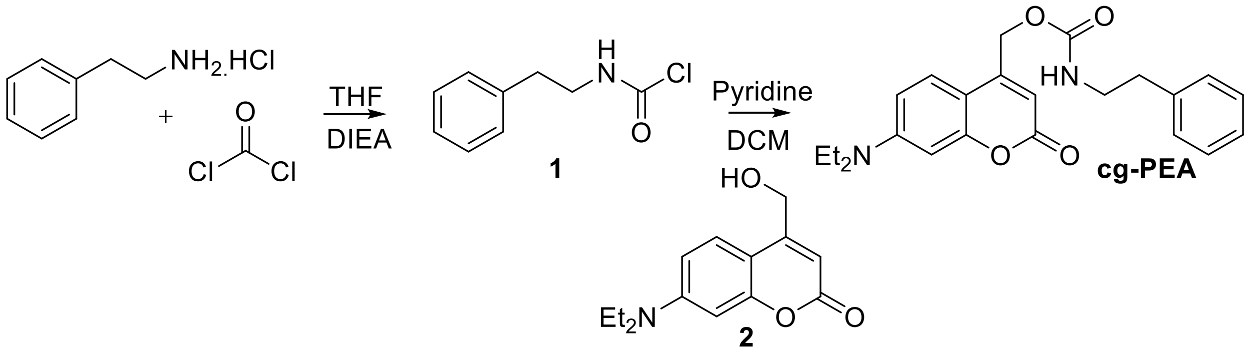


Intermediate **1** was prepared from a solution of phenylethylamine hydrochloride (500 mg, 3.17 mmol) in dry THF (50 mL), treated with DIEA (413.6 mg, 307 µL, 3.2 mmol), and stirred for 5 min at 0 °C. After dropwise addition of phosgene (4.3 mL, 10 mmol), the reaction mixture was stirred for an additional 90 min at 0 °C. The reaction mixture was transferred onto a mixture of EtOAc and $H_2O$ (1:1, 200 mL), the layers were separated, the organic layer washed with $H_2O$ (1 x 50 mL) and sat. NaCl solution (1 x 50 mL) and dried over $Na_2SO_4$. The solvent was removed under reduced pressure and the N-[ethylbenzyl]chloroformamide **1** was engaged in the next step without further purifications.

7-diethylamino-4-hydroxymethylcoumarin **2** were synthesized by reproducing the methodology reported by Walter et al. [1]. In a separated flask, a solution of 7-diethylamino-4-hydroxymethylcoumarin **2** (500 mg, 2.04 mmol) in a mixture of DCM and pyridine (3:1, 20 mL) was stirred for 5 min at 0 °C. Intermediate **1** was dissolved in DCM (2 mL) and this solution was added dropwise to the reaction mixture. The reaction was protected from light and allowed to reach r.t. overnight. The reaction mixture was transferred onto a mixture of DCM and $H_2O$ (1:1, 100 mL), the layers were separated, the organic layer washed with $H_2O$ (2 x 50mL) and sat. NaCl solution (1 x 50 mL) and dried over $Na_2SO_4$. The solvent was removed under reduced pressure and the crude purified by flash chromatography (eluent: cyclohexane/EtOAc 3:1). The target compound was obtained as a pale brown powder (350 mg, 0.8 mmol, yield= 39%).

1H-NMR (400 MHz, CDCl3) δ = 7.26-7.13 (m, 6H), 6.51 (d, 1H), 6.44 (s, 1H), 6.03 (s, 1H), 5.15 (s, 2H), 4.92 (s, 1H), 3.46 (s, 2H), 3.35 (q, 4H), 2.79 (s, 2H), 1.14 (t, 6H).

13C-NMR (100 MHz, CDCl3) δ = 12.44, 19.35, 36.01, 42.38, 44.75, 47.53, 61.72, 97.82, 106.04, 106.09, 108.64, 124.40, 126.65, 128.79, 138.49, 150.39, 150.63, 155.44, 162.04 ppm.

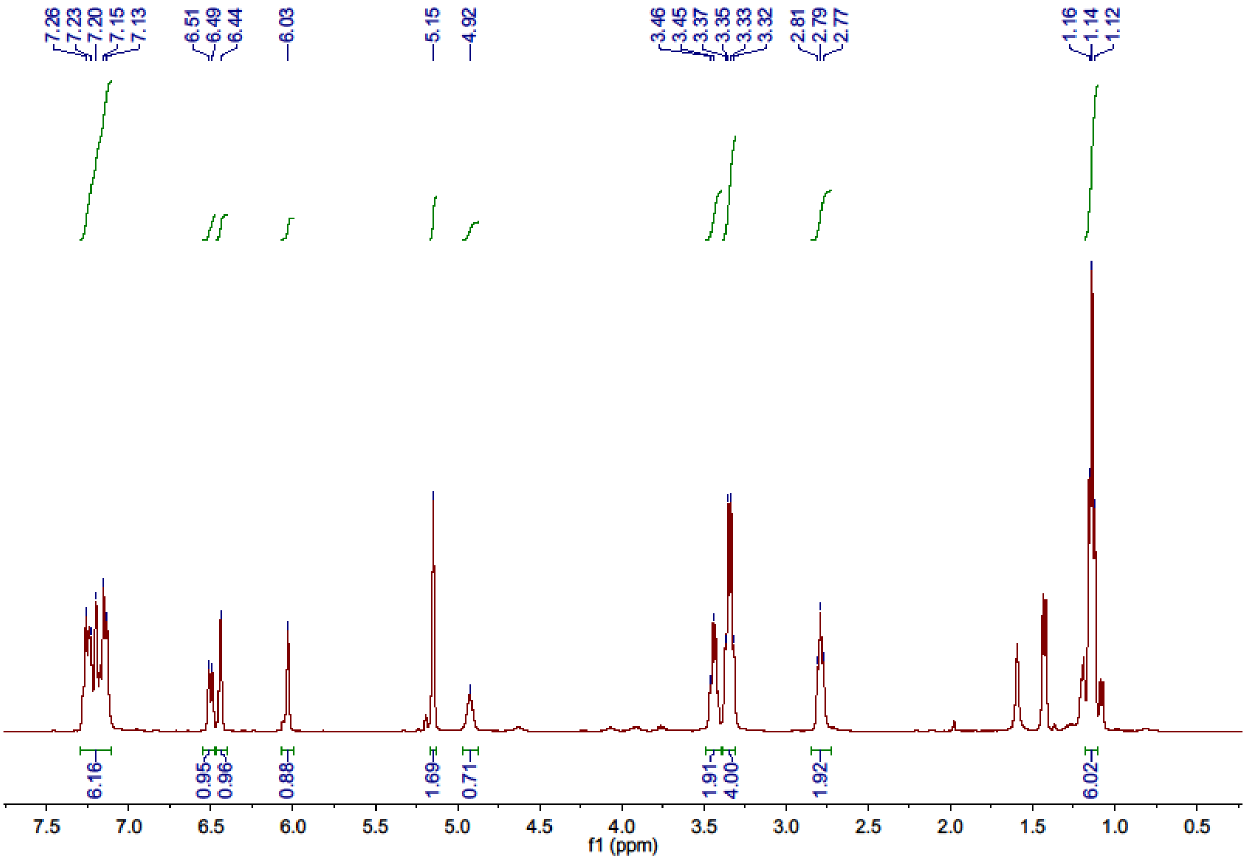


$^1$H-NMR spectrum of cg-PEA

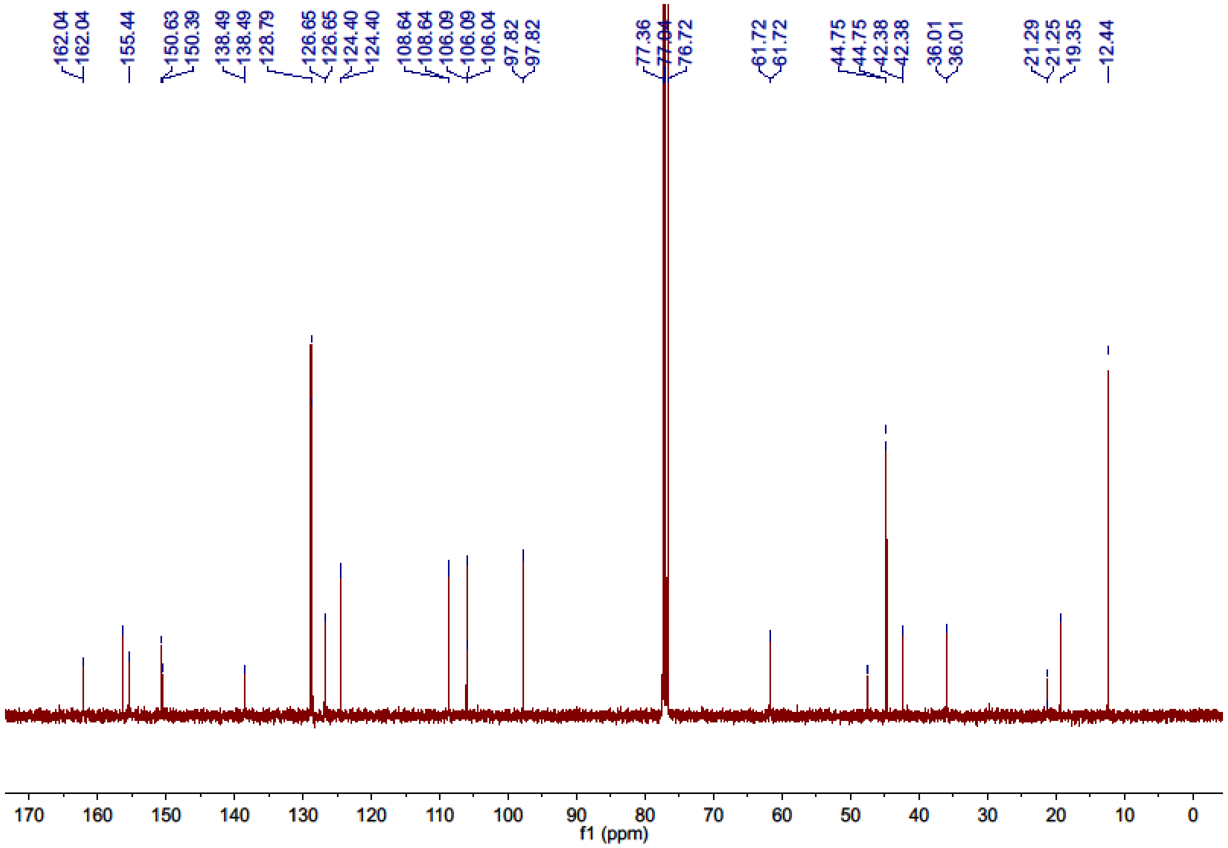


$^{13}$C-NMR spectrum of cg-PEA
